# Dynein-2 is tuned for the A-tubule of the ciliary doublet through tubulin tyrosination

**DOI:** 10.1101/2025.01.27.635025

**Authors:** Haoqiang K. He, Shintaroh Kubo, Qianru H. Lv, Azusa Kage, Muneyoshi Ichikawa

## Abstract

Eukaryotic cilia and flagella are thin structures present on the surface of cells, playing vital roles in signaling and cellular motion. Cilia structures rely on intraflagellar transport (IFT), which involves dynein-2 for retrograde and kinesin-2 for anterograde movements along doublet microtubules. Unlike dynein-1, which works on singlet microtubules within the cytoplasm, dynein-2 specifically works on the doublet microtubules inside the cilia. Previous cryo-electron tomography studies have shown that retrograde IFT, driven by dynein-2, occurs on the A-tubule of the doublet, suggesting a specialized regulatory mechanism involving dynein-2. However, the molecular basis of this specificity remains unclear. Here, we investigated this mechanism using cryo-electron tomography (cryo-ET) with Volta Phase Plate (VPP), molecular dynamics (MD) simulations, and biochemical analysis. Our biochemical assay revealed that the microtubule-binding domain of dynein-2 exhibits a higher affinity for the ciliary doublet microtubule compared to dynein-1. Cryo-ET with VPP further visualized the preferential binding of dynein-2 to the A-tubule of the doublet microtubule. MD simulations suggest that the preferential binding of dynein-2 is attributed to the tyrosinated tubulin in the A-tubule. These findings uncover a tyrosination-dependent regulatory mechanism that governs the bidirectional transport of IFT on doublet microtubules, providing new insights into the spatial and functional specialization of ciliary transport systems.

## Introduction

In eukaryotic cells, cilia (also referred to as flagella) are highly conserved hair-like organelles that extend from the cell surface and have a wide range of functions. Non-motile primary cilia play roles in signaling, and motile cilia provide motility for specific cells like sperm. Both types of cilia possess nine doublet microtubules (doublets) composed of a complete A-tubule and an incomplete B-tubule (Gibbons, 1981; Ishikawa *et al*., 2011). The ciliary doublets are covered with various microtubule-associated proteins (MAPs) for stabilization, as well as protein complexes involved in ciliary beating, such as axonemal dyneins, radial spokes, and the Nexin-dynein regulatory complex, except for the area facing the membrane (Walton *et al*., 2023; Leung *et al*., 2025). This empty space is used as a railway for intraflagellar transport (IFT) (Fig 1). The IFT is a cargo transport process that takes place in the space between doublets and ciliary membranes and plays a vital role in the assembly and maintenance of the complex structure of cilia (Webb *et al*., 2020). The IFT complex consists of two sub-complexes, IFT-A and IFT-B, and IFT proteins are linked to various human ciliopathies (Reiter *et al*., 2017). These sub-complexes assemble sequentially to form IFT trains, which are transported along doublets by the motor proteins dynein-2 and kinesin-2.

**Figure 1.**
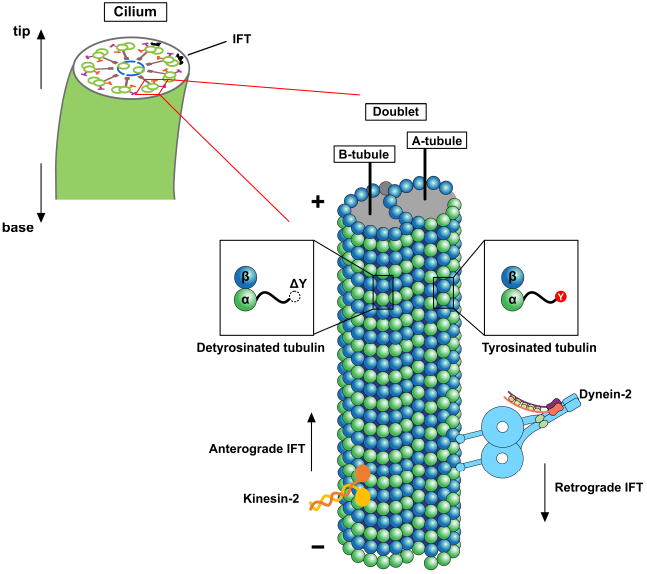
Schematic illustration of the structure of the cilia and the mechanism of the IFT. Dynein-2 drives retrograde IFT toward the ciliary base on A-tubules, while kinesin-2 powers anterograde IFT toward the ciliary tip on B-tubules. In the doublets, A-tubules are enriched with tyrosinated tubulin, whereas B-tubules are enriched with detyrosinated tubulin. Microtubule polarity is indicated by (+) and (−).

Dynein-2 is a multiprotein complex comprising two heavy chains (HCs), a heterodimer of intermediate chains (ICs) (WDR60/WDR34), two light intermediate chain-3 (LIC3), homodimeric light chain (LC) DYNLRB, three homodimeric LC8, and heterodimeric LC (TCTEX/TCTEX1D2) (Toropova, *et al*., 2019) (Fig 2A). Dynein-2 and dynein-1 share a similar domain organization, including a tail domain and a head domain, which is composed of the AAA+ motor domain, coiled-coil stalk, and microtubule-binding domain (MTBD). Dyneins interact with microtubule (MT) with MTBD that undergoes the transition of MT low-affinity and MT high-affinity states (Gibbons *et al*., 2005; Redwine *et al*., 2012). Despite the similarity, their functions are distinct: dynein-1 operates on singlet MTs (singlets) in the cytoplasm, whereas dynein-2 is specialized for transportation on ciliary doublets (Yildiz & Zhao, 2023).

**Figure 2.**
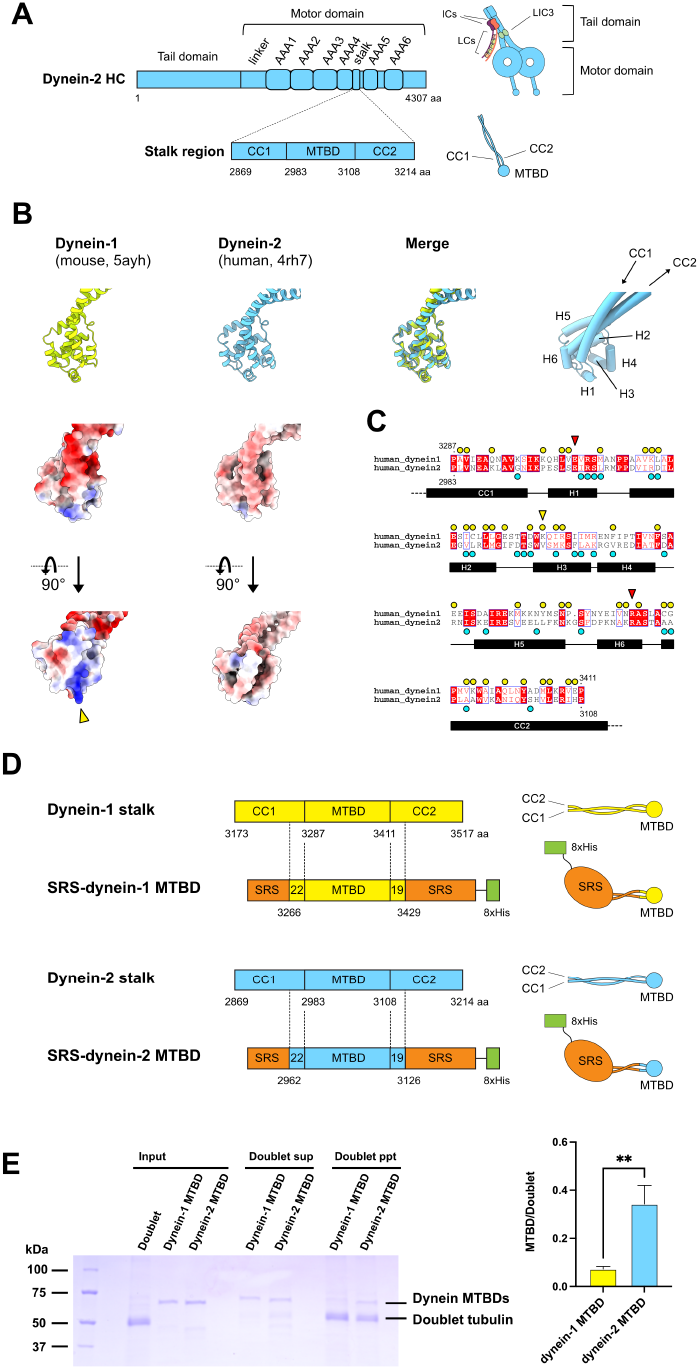
Comparison of MTBDs of dynein-1 and dynein-2. **(A)** Schematic representation of the dynein-2 HC and stalk region and their structures. The HC includes the tail domain and the motor domain. The stalk region of the motor domain is highlighted, and CC1, MTBD, and CC2 within the stalk region are indicated. The diagram includes subunits associated with the dynein-2 tail domain. **(B)** Comparison of the dynein-1 MTBD and dynein-2 MTBD structures. Dynein-1 stalk structure (top row, leftmost, PDB ID: 5ayh) (Nishikawa *et al*., 2016) and dynein-2 motor domain structure (top row, second, PDB ID: 4rh7) (Schmidt *et al*., 2014) are aligned based on the MTBD region. Matched structures (top row, third) show that the general structures of MTBD of dynein-1 and dynein-2 are similar. Surface models of dynein-1 and dynein-2 MTBDs (second row), colored to show electrostatic potential ranging from red (−10 kcal/(mol·*e*)) to blue (+10 kcal/(mol·*e*)). 90° rotated views of surface models (bottom row) show that the contact surfaces of dynein-1 and dynein-2 MTBDs to the tubulin lattice are distinct. Helix numbers, CC1, and CC2 are indicated in the rightmost panel. Yellow arrowhead indicates K3336 residue of H3 from dynein-1 (residue number is based on human dynein-1). **(C)** Sequence alignment of human dynein-1 and human dynein-2 MTBD regions. Amino acid sequences of dynein-1 MTBD region (NCBI Reference Sequence: NP_001367.2) and dynein-2 MTBD region (NCBI Reference Sequence: NP_001368.2) are aligned. The alignment was performed using ClustalW (Thompson, 1994), and the figure was generated using ESPript 3.0 (Robert & Gouet, 2014). Note that the sequence of mouse dynein-1 MTBD region (PDB ID: 5ayh) and human dynein-1 MTBD region are exactly the same. The yellow circles indicate the amino acid residues conserved between dynein-1, and the cyan circles show the amino acid residues conserved between dynein-2, respectively (see also Fig S1). K3336 of dynein-1 is shown by the yellow arrowhead. Red arrowheads indicate the residues previously shown as important in MT binding. Positions of CC1, CC2, and helices are indicated at the bottom. **(D)** Schematic diagrams of the stalk regions and the MTBD constructs used in this study. CC1 and CC2 of the MTBD regions were merged with SRS coiled-coil (orange) to fix MTBD structures to MT strong binding conformations, and 8×His-tag (green) was added for purification. **(E)** Copelleting assay of dynein MTBD constructs with doublets. MTBD constructs were incubated with doublets, ultracentrifuged, and analyzed by SDS-PAGE (left), and the bands were quantified (right). The band of the dynein-2 MTBD construct in precipitation (ppt) fraction was more prominent compared with that of dynein-1 MTBD construct. Error bars represent the SD from three independent experiments. Statistical analyses were performed using an unpaired *t* test (\*\**P*≦0.01).

In the IFT process, the anterograde IFT train driven by kinesin-2 transports cargo from the cytoplasm to the tip of the cilia. Then, the anterograde IFT train remodels into the retrograde IFT train and transports back toward the cell body by dynein-2 (Lacey *et al*., 2024) (Fig 1). To avoid the collision of anterograde and retrograde IFT trains in the limited space between the doublet and membrane, the doublet serves as dual railway tracks for IFT. From the previous cryo-electron tomography (cryo-ET) study of flagella, anterograde IFT trains were found on the B-tubule, while retrograde IFT trains were found on the A-tubule (Stepanek & Pigino, 2016).

Since the tubulins composing A- and B-tubules have different post-translational modifications (PTMs) like tyrosination/detyrosination, polyglutamylation, and polyglycylation (Westermann & Weber, 2003), the different PTMs could lead to the sorting of the anterograde and retrograde IFTs. A recent study has shown that the polyglycylation is present both in A- and B-tubules (Alvarez Viar *et al*., 2024) and, therefore, cannot be the cause for sorting the IFTs. The same study located the polyglutamylation on the protofilament (PF)-9 of the B-tubule that is not the track of IFT.

The tyrosination/detyrosination occurs on the C-terminus of the α-tubulin (Sanyal *et al*., 2023). In most cases, expressed α-tubulin has tyrosine residue at the C-terminus (tyrosinated), and this tyrosine residue can be enzymatically removed by tubulin carboxypeptidases (TCPs) (detyrosination). The removal of the C-terminal tyrosine residue exposes the glutamic acid residue at the C-terminus. A tyrosine residue can be re-attached to the detyrosinated tubulin by the tubulin tyrosine ligase (TTL). The A-tubule is shown to be enriched with tyrosinated tubulins, and the B-tubule has detyrosinated tubulins (Johnson, 1998) (Fig 1). Very recently, the tyrosination/detyrosination balance of the doublet was shown to be important for the proper sorting of IFT trains (Chhatre *et al*., 2024). Despite these advances, the exact mechanism by which retrograde IFT is selectively recruited to the A-tubule remains unidentified.

Here, we used cryo-ET analysis aided by Volta Phase Plate (VPP) of *in vitro* reconstitution system, molecular dynamics (MD) simulations, and biochemical analysis, and provide evidence that dynein-2 exhibits distinct binding properties to doublets compared to dynein-1, and dynein-2 prefers to bind to the A-tubule of the doublet due to its tyrosinated state.

## Results

### Dynein-2 is tuned for the binding to the ciliary doublet

We hypothesize that the MT-binding property of dynein-2 differs from those of dynein-1, reflecting the distinct intracellular functional locations. To verify this, we compared available MTBD structures and sequences of dynein-2 and dynein-1 (Fig 2B and C). The overall structures of the MTBDs were highly similar, with an RMSD of 1.1 Å for the MTBD region. Some residues of the *Dictyostelium* dynein that were previously shown to form salt bridges with the tubulin residues (Uchimura *et al*., 2015) were conserved in human dynein-2 and dynein-1 HCs. For instance, E3002 of H1 from dynein-2 (equivalent to E3306 of dynein-1 and E3390 from *Dictyostelium* dynein) and R3081 of H6 from dynein-2 (equivalent to R3384 of dynein-1 and R3469 from *Dictyostelium* dynein) (Fig 2C, red arrowheads). Despite these similarities, the surface charge of the MTBD in dynein-2 was notably distinct from that in dynein-1 (Fig 2B). A key difference was the absence of K3336 from H3 of dynein-1 in the dynein-2 MTBD (Fig 2B and C, yellow arrowheads). Other residues conserved within dynein-2 MTBD were also distinct from those in dynein-1 MTBD (Fig 2C and Fig S1).

To further explore these findings, we performed an MT co-pelleting assay using MTBD constructs and native doublets. We used the MTBD construct fused with seryl-tRNA synthase (SRS) for dynein-2 and dynein-1, respectively (Fig 2D). The constructs used in this study are equivalent to the α registry (MT-high affinity) 22:19 construct in (Carter *et al*., 2008). For doublets, we purified the native doublets from cilia and removed the associated proteins (Fig S2). Our previous structural analyses have shown that doublets containing Microtubule Inner Proteins (MIPs) but without MAPs outside can be prepared with this procedure (Ichikawa *et al*., 2017; Ichikawa *et al*., 2019). After incubating dynein MTBD constructs with doublets, the bound dynein MTBD proteins were pelleted with doublets by centrifugation and analyzed by SDS-PAGE. As a result, dynein-2 exhibited higher binding compared with dynein-1 (Fig 2E) consistent with dynein-2’s function on doublets.

### Dynein-2 is tuned for the binding to the A-tubules

To further explore how dynein-2 interacts with the doublet, we exploited an *in vitro* reconstitution system constituted of minimal components. GST-Dyn2, lacking a tail domain and having mutations to mimic active dynein-2 with two heads open (Tropova *et al*., 2017) (Fig 3A), was incubated with native doublets with clean outer surfaces as purified above (Fig 3B). After the centrifugation, GST-Dyn2 was detected in the pellet fraction together with the doublets, and as the concentration of GST-Dyn2 increased, the binding was saturated (Fig 3C).

**Figure 3.**
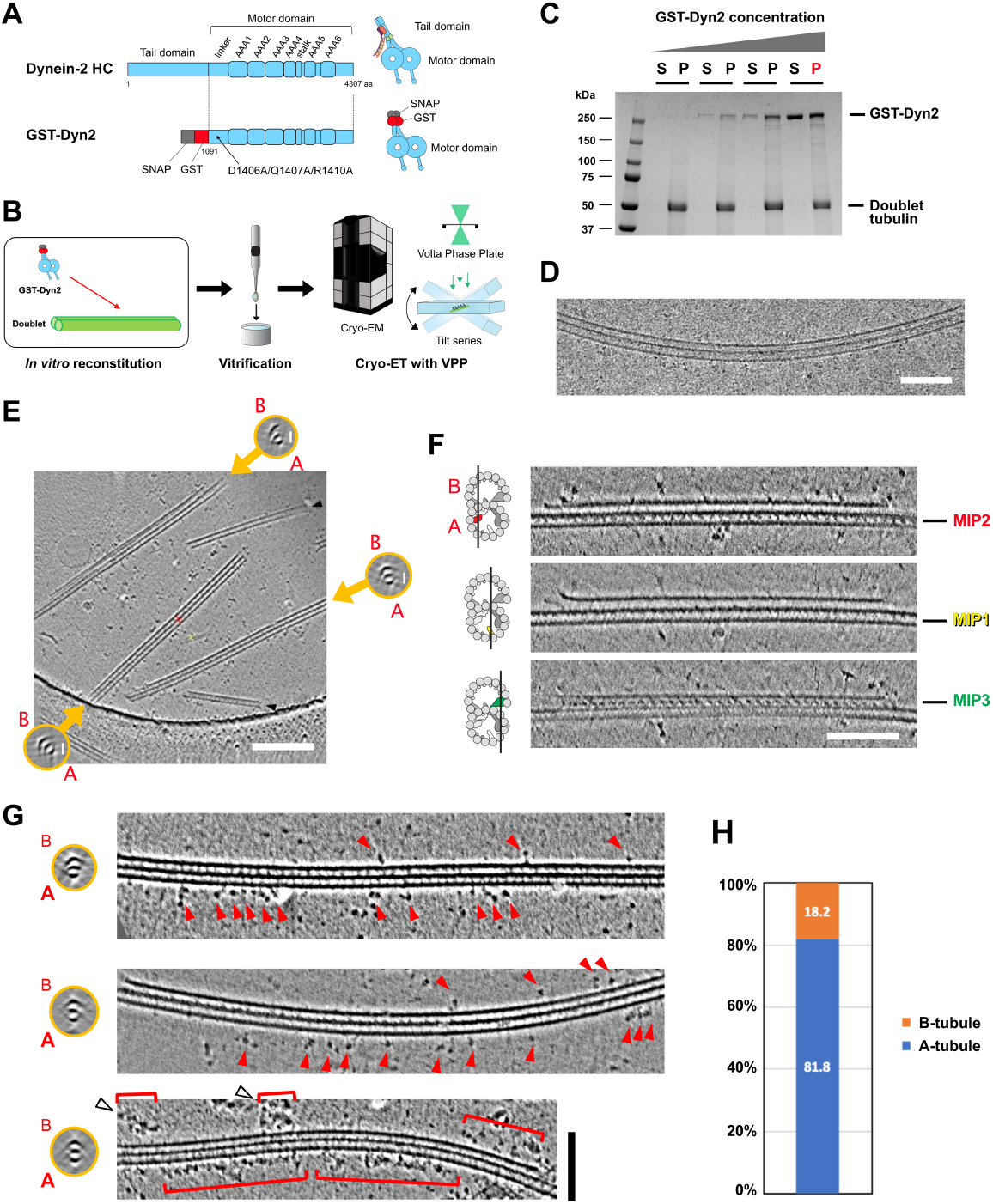
Cryo-ET analysis of GST-Dyn2 bound to doublets. **(A)** Schematic illustrations of the dynein-2 HC and GST-Dyn2 construct. The tail domain of the dynein-2 HC was truncated, and the GST-tag (red) was added for dimerization with the SNAP-tag (grey). Mutations (D1406A/Q1407A/R1410A) were introduced to separate two motor domains. **(B)** The analysis workflow of the binding mode of the GST-Dyn2 to doublets. GST-Dyn2 was incubated with doublets and vitrified. The grid was loaded into cryo-EM, and tilt series were acquired using the VPP. **(C)** The SDS-PAGE image of the co-pelleting assay of GST-Dyn2 and doublets. A fixed concentration of doublets (500 μg/ml) was incubated with increasing concentrations of GST-Dyn2 (0, 112.5, 225, and 450 μg/ml), centrifuged, and the supernatant fraction (S) and precipitation fraction (P) were analyzed by SDS-PAGE. The precipitated sample (450 μg/ml, indicated in red) was resuspended in a buffer and used for vitrification. **(D)** A typical cryo-EM image of GST-Dyn2 decorated doublet. The image was taken with a single tilt with a nominal 2 μm defocus. There are more GST-Dyn2 molecules bound to one side of the doublet. Scale bar, 100 nm. **(E)** Example of a tomographic slice of reconstructed GST-Dyn2 decorated doublets. Cross-sectional views are shown for each doublet in orange circles, and A- and B-tubules are indicated. Scale bar, 200 nm. Note that there are several singlets due to sample preparations. See also Movie S1 and S2. **(F)** Longitudinal tomographic slice of the representative reconstructed doublets’ 3D structures. Black lines in the illustrations of the doublet on the left indicate the sections on the right panels. MIP structures distinctive for A- or B-tubules are highlighted in the model and indicated in the tomographic slices. Naming and colorings are adopted from (Ichikawa *et al*., 2017; Ichikawa *et al*., 2019). Scale bar, 100 nm. **(G)** Representative examples of tomograms of GST-Dyn2 bound predominantly on the A-tubules of the doublets. Cross-sectional view (left, orange circles) and longitudinal views (right) are shown for each doublet, and A- and B-tubules are indicated. All doublet structures are shown in the figure so that the A-tubule sides face the bottom. Red arrowheads indicate the dynein2 motor domains. Red brackets indicate the clustered GST-Dyn2 on the doublets. White arrowheads indicate the GST-Dyn2 clusters detaching from doublets. Scale bar, 100 nm. **(H)** Quantification result showing that GST-Dyn2 tends to bind to the A-tubules side of the doublets. Dynein-2 motor domains were counted and assessed if the A- or B-tubule has more dynein2 motor domains. 81.8% of doublets (18 out of 22) have more dynein-2 motor domains on the A-tubule side.

The sample was vitrified and observed with the saturated condition by cryo-electron microscopy (cryo-EM). Because of the preferred orientation, the doublets were primarily observed in side views, with both tubules visible. From the obtained single tilt cryo-EM image, GST-Dyn2 molecules were accumulated on one side of the doublet (Fig 3D). To further identify whether GST-Dyn2 binds more on the A- or B-tubule sides, we performed cryo-ET analysis using the Volta Phase Plate (VPP) (Fukuda *et al*., 2017; Imhof *et al*., 2020; Pöge *et al*., 2021) to enhance contrast and visualize the dynein-2 motor domains clearly (Fig 3B and Movie S1 and S2). In the reconstructed tomograms, the A- and B-tubules of the doublets were readily distinguished both by the morphology of the tubules and the MIPs distinctive for the A- or B-tubule (Fig 3E and F).

By identifying the A- and B-tubules of the doublet, GST-Dyn2 molecules were found predominantly on the A-tubule side without external components (Fig 3G). By quantifying the numbers of the dynein-2 motor heads, the majority of doublets (81.8%, 18 out of 22) have more dynein-2 on the A-tubule side (Fig 3H). Furthermore, compared with the continued clustering of the GST-Dyn2 on the A-tubule, the clustering of the GST-Dyn2 found on the B-tubule were shorter and often detaching from the tubule (Fig 3G, white arrowheads).

### Dynein-2 prefers tyrosinated tubulin lattice

Previously, it has been shown that the A-tubule of the doublet is enriched with tyrosinated tubulins while the tubulins forming the B-tubule are detyrosinated (Johnson, 1998). To test if the preferential binding of the GST-Dyn2 to the A-tubule observed in our cryo-ET was due to the tyrosination/detyrosination states of the doublet tubulin lattice, we performed MD simulations.

Two dynein-2 heads, one with an MT low-affinity state and the other with an MT high-affinity state, representing a dynein-2 dimer, were placed on the tubulin lattice with or without tyrosination (Fig 4A and S3A). To observe the initial diffusional motion of the low-affinity state MTBD right after it shifts from the high-affinity state to the low-affinity state, the low-affinity state MTBD was placed at the MT binding site (see Materials & Methods for details). Three relative positions of the two heads were used as initial positions, and the trajectories of the low-affinity state dynein-2 head were simulated by coarse-grained MD (Fig 4B). After the calculation, the probabilities of the existence of low-affinity MTBD were plotted for both tyrosinated and detyrosinated tubulin lattice, respectively (Fig 4C and D, Fig S3B and C). The heat maps were generated for each initial position, and the sum of three initial positions were plotted. In the sum result, dynein-2 MTBD tended to stay on the same position (1,1) in the tyrosinated tubulin lattice compared with the detyrosinated tubulin lattice (Fig 4C and D). Furthermore, overflow proportion, which include the detachment from the tubulin lattice, was higher in detyrosinated tubulin lattice (20.09%) compared with tyrosinated tubulin lattice (10.70%), suggesting that dynein-2 molecules are easier to detach from the detyrosinated tubulin lattice. This is consistent with our cryo-ET observation showing that GST-Dyn2 molecules detach from the B-tubule.

**Figure 4.**
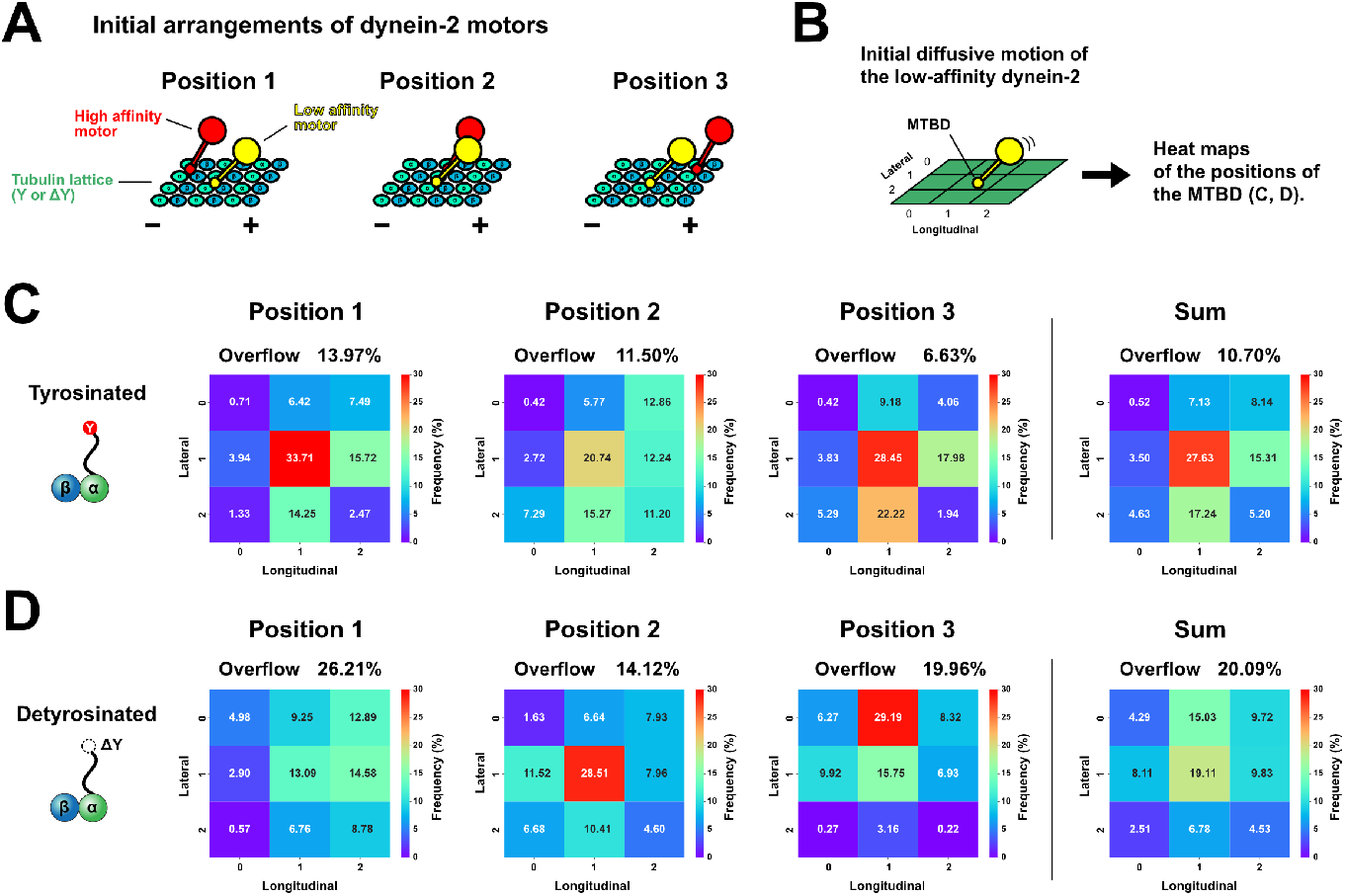
MD simulation results of dynein2 dimer on tyrosinated or de-tyrosinated tubulin lattice. **(A)** Illustrations showing the setup of MD simulations. The low-affinity dynein-2 motor domain (yellow) and the high-affinity dynein-2 motor domain (red) were placed on the neighboring protofilament (PF) with three arrangements (positions 1, 2, and 3). Either tyrosinated (Y) or detyrosinated (ΔY) tubulin lattice was used. MT polarities are indicated. **(B)** A schematic of trajectory analysis of the low-affinity motor. The initial diffusional motion of the low-affinity dynein-2 motor was simulated, and the positions of MTBD were plotted in a 3×3 area in (C and D). **(C and D)** Heat maps of the low-affinity dynein-2 MTBD position after the MD simulations with tyrosinated tubulin in B and detyrosinated tubulin in C. The mass center-of-gravity coordinates of the MTBDs of the low-affinity dynein-2 obtained from their trajectories are shown in the heat map (red: higher frequency; blue: lower frequency). The sum of the positions 1, 2, and 3 are shown in the rightmost panel. The center (1, 1) is the initial position of low-affinity dynein-2 MTBDs. Overflow refers to the cases where MTBDs diffused outside the 3×3 area including detachment. See also Fig S3.

Furthermore, in the result from initial position 3 with detyrosination, the low-affinity leading head was observed to block the path of the high-affinity trailing head (Fig S3D). When the trailing head transitions to the low-affinity state, the dynein-2 dimer will not move toward the minus end since the next binding site on the same PF is unavailable. This type of blockage was not observed for the tyrosinated tubulin lattice.

## Discussion

Here, we showed that dynein-2 exhibited distinct binding properties compared to dynein-1, showing a higher affinity for ciliary doublets. Our cryo-ET analysis showed that dynein-2 preferentially binds to the A-tubule of the doublet, and our MD simulations revealed that this preference is likely due to reduced affinity for the detyrosinated B-tubule. In the anterograde IFT process, dynein-2 takes stacked conformation within IFT train cargo as transported by kinesin-2 (Jordan *et al*., 2018; Toropova *et al*., 2017). At the ciliary tip, dynein-2 is released from the anterograde IFT train and activated by opening two heads. Based on our results, these active dynein-2 molecules can bind to doublets on either A- or B-tubule, but the dynein-2 molecules will be easily detached from the B-tubule side, and dynein-2 molecules on the A-tubule side start the retrograde IFT (Fig 5). Recently, it was shown that affinities of IFT trains to the A- or B-tubules were affected by the level of tyrosination/detyrosination of the doublets (Chhatre *et al*., 2024). Still, the molecular mechanism underlying was not revealed. Our work shows that dynein-2 recognizes the doublet’s tyrosination/detyrosination states for the retrograde IFT trains.

**Figure 5.**
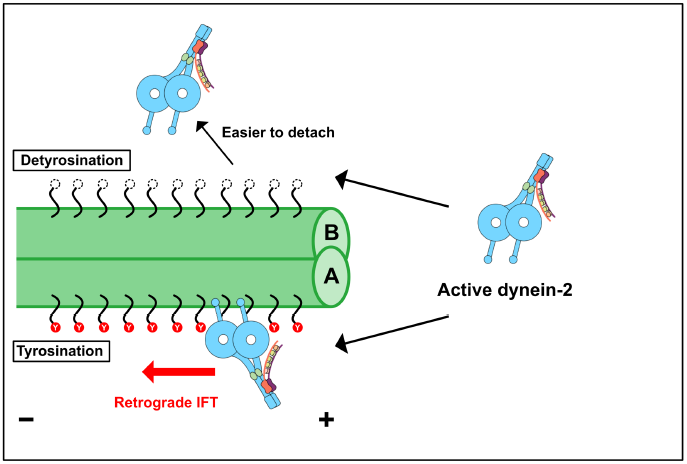
Model of the track recognition mechanism of dynein-2. At the tip of the cilia, dynein-2 will be released from anterograde IFT driven by kinesin-2. Released dynein-2 molecules get activated in unelucidated mechanisms by opening two motor domains. Active open dynein2 molecules bind to the doublets on either the A- or B-tubule side. Since the tubulin lattice from the B-tubule is detyrosinated, dynein-2 molecules are easier to detach. Dynein-2 molecules bound to the A-tubule tend to stay on the tyrosinated tubulin lattice, and these dynein-2 molecules exhibit unidirectional motion and power retrograde IFT.

Dynein-2 has a distinct surface charge compared with dynein-1 (Fig 2B), most likely for selecting its pathway within the doublets. In other types of dynein, like dynein-c (DHC9), flap structures are inserted between H2 and H3 of the MTBD (Kato *et al*., 2014). For γHC (ODA2) of outer arm dynein, LC1 is associated with the MTBD region (Ichikawa *et al*., 2015; Toda *et al*., 2020). These flap structures and LC1 are thought to be important for the interaction with doublets area where two PFs have wider spaces (Lacey *et al*., 2019; Rao *et al*., 2021). Dynein-2 does not use flap or associated protein but has changed its amino acid sequences to adapt to the rail. Previously, H3 of dynein-1 MTBD was shown to play a role in the interaction with MT (Redwine *et al*., 2014). The absence of a positively charged residue in the H3 region of dynein-2 might contribute to its preference for tyrosinated tubulin because the glutamic acid residue is exposed at the C-terminus of the detyrosinated tubulin.

Our analyses show that dynein-2 prefers tyrosinated tubulin lattice over detyrosinated tubulin lattice. For kinesin-2, which drives the anterograde IFT, previous research has shown that the detyrosinated tubulin lattice enhances both its processivity and velocity (Sirajuddin *et al*., 2014). This difference in the sensitivities to the tyrosination state of dynein-2 and kinesin-2 is crucial for sorting the retrograde and anterograde IFT. The cytoplasmic dynein from yeast, which lacks cilia, is not affected by the tyrosination states of the MT (Sirajuddin *et al*., 2014). *In vitro* study has shown that kinesin-1 itself is not affected by tyrosination state (Sirajuddin *et al*., 2014). Therefore, in the species having cilia, motors involved in IFT and doublets have co-evolved so that the doublets provide specific trajectories for IFTs of opposing directions. Similar specialization of MTs by tyrosination/detyrosination states has also been observed in singlets from cytoplasm. As for the cytoplasmic dynein-1, while mammalian dynein-1 itself is insensitive to the tyrosination states of the MT (McKenney *et al*., 2016), mammalian dynein-1 has evolved its binding partners like CLIP-170 and p150Glued of dynactin complex to sense the tyrosination state of the singlet (McKenney *et al*., 2016; Nirschl *et al*., 2016). Dynein-2, which does not have a modulator like CLIP-170 or dynactin, has likely tuned its MTBD to bind to tyrosinated tubulin lattice through evolution. Several kinds of kinesins are known to be sensitive to tyrosination/detyrosination states of singlets. For instance, kinesin-3 and kinesin-5 preferentially walk on tyrosinated singlets (Tas *et al*., 2017; Kahn *et al*., 2015). Based on these, tubulin tyrosination/detyrosination is a conserved strategy for sorting the motors between singlets and doublets.

Our results suggest that the MTBD of dynein-2 is fine-tuned for binding to the A-tubule. However, identifying the exact residues critical for track recognition will require high-resolution structural analyses in future studies. Additionally, the mechanism by which dynein-2 is activated after being released from anterograde IFT remains unclear. Furthermore, while it is known that PTMs exhibit distinct patterns on doublets, the processes that generate these patterns have yet to be elucidated.

In conclusion, we identified dynein-2 as a key component of the track recognition mechanism underlying retrograde IFT, an essential process for ciliary assembly, and one associated with human ciliopathies.

## Materials and Methods

### Purification of MTBD constructs

The sequences coding human dynein-1 MTBD and part of coiled-coil region (NCBI Reference Sequence: NP_001367.2, 3266-3429 aa) or human dynein-2 MTBD and part of coiled-coil region (NCBI Reference Sequence: NP_001368.2, 2962-3126 aa) were inserted into modified pET-42a vector, respectively. The plasmids were transformed into *E. coli* BL21(DE3) cells and the cells were cultured at 37°C in LB media containing 30 µg/ml kanamycin until the OD_600_ reached around 0.6. Protein expression was induced by adding 0.4 mM isopropyl-1-thio-β-D-galactopyranoside and incubated at 20°C for 3 hours. From here, the procedure was performed at 4°C or on ice. Cells were harvested by centrifugation and were resuspended in lysis buffer (50 mM Tris-HCl pH 8.0, 150 mM NaCl, 20 mM imidazole) supplemented with 1 mM phenylmethylsulfonyl fluoride (PMSF) and cOmplete™ EDTA-free Protease Inhibitor Cocktail (Roche). The cells were homogenized, and the lysate was centrifuged at 15,000g for 30 minutes using a SORVALL LYNX 6000 Centrifuge (Thermo Fisher Scientific) and an A27-8×50 rotor (Thermo Fisher Scientific). The supernatant was retrieved and incubated with Ni-NTA agarose (30250, QIAGEN) with rotation for 1 hour. The column was washed with wash buffer (50 mM Tris-HCl pH 8.0, 250 mM NaCl, 20 mM imidazole), and bound proteins were eluted with elution buffer (50 mM Tris-HCl pH 8.0, 150 mM NaCl, 300 mM imidazole). The buffer of the elution fraction was exchanged for NAP5 buffer (50 mM Tris-HCl pH 8.0, 100 mM NaCl) by NAP™-5 Columns (17-853-02). The obtained proteins were frozen with liquid nitrogen and stored at −80°C.

### Purification of GST-Dyn2

GST-Dyn2 construct with mutations (D1406A/Q1407A/R1410A) was expressed and purified as in (Toropova *et al*., 2017). pFastBac vector containing ZZ-tag, TEV cleavage site, SNAPf-tag, GST tag, and dynein-2 heavy chain sequence (1091-4307 aa, codon-optimized) between Tn7 sites was transformed to DH10EMBacY cells. Colonies with EMBacY bacmid containing GST-Dyn2 cassette between Tn7 sites were chosen by blue/white selection, cultured and bacmids were purified. Sf9 cells (Addgene, 64064) were cultured in Insect-XPRESS Medium supplemented with L-glutamine at 27°C with shaking at 100 rpm. Sf9 cells were transfected with bacmids using FuGene HD transfection reagent (Promega), and transfection efficiency was checked by YFP fluorescence from EMBacY. The media (V0) was collected and added to a 50 ml Sf9 culture. After incubation for 3 days, the cells were removed from the media by centrifugation, and the media (V1) was collected and stored at 4°C. 2.5 ml of V1 was then added to 250 ml Sf9 culture. After three days, cells were harvested by centrifugation, washed with 1×PBS, flash-frozen in liquid nitrogen, and kept at −80°C until purification. For purification, cells were resuspended in 20 ml purification buffer (30 mM 4-(2-hydroxyethyl)-1-piperazineethanesulfonic acid [HEPES] pH 7.4, 300 mM KCl, 50 mM potassium acetate, 2 mM magnesium acetate, 1 mM ethylene glycol tetraacetic acid [EGTA], 10% glycerol, 1 mM dithiothreitol [DTT], 0.2 mM adenosine triphosphate [ATP], 1 mM PMSF) containing cOmplete™ EDTA-free Protease Inhibitor Cocktail (Roche). Cells were lysed and ultracentrifuged (Type 70 Ti rotor, 183,960g, 30 min, 4°C), and the supernatant was incubated with 1 ml of IgG Sepharose 6 resin (GE Healthcare) pre-equilibrated with purification buffer. The resin was collected by centrifugation (670g, 5 min 4°C) and washed with 20 ml of purification buffer twice and 20 ml of TEV buffer (50 mM Tris pH 7.5, 150 mM potassium acetate, 2 mM magnesium acetate, 1 mM EGTA, 10% glycerol, 1 mM DTT, 0.2 mM ATP). The dynein-2 construct with SNAPf-tag and GST tag was removed from the resin by incubating with 4 ml of TEV buffer containing 100 μg TEV protease overnight at 4°C with rotating. The eluted dynein-2 construct was concentrated using Amicon Ultra Centrifugal Filter Unit (100 kDa cutoff, Millipore), and ultracentrifuged (TLA 110 rotor, 337,932g, 6 min), and the supernatant was flash-frozen in liquid nitrogen and stored at −80°C.

### Purification of doublets with clean outer surface

Native doublets with clean outer surfaces were prepared from *Tetrahymena thermophila* as described in (Black *et al*., 2021). *Tetrahymena* cells (SB255 strain) were cultured in 1 L of SPP media (1% proteose peptone No. 3, 0.2% glucose, 0.1% yeast extract, 0.003% ethylenediaminetetraacetic acid iron (III) sodium salt [Fe-EDTA]) until the OD_600_ reached 0.7. Cells were harvested by low-speed centrifuge (700g, 10 min, 4°C) and concentrated to 25 ml with SPP media. The procedure was performed at 4°C hereafter otherwise noted. Cilia were removed from the cell bodies by adding the final 1 mg/ml dibucaine and swirling for 1 min. The cell bodies were removed by slow-speed centrifugation (2,000g, 7 min, 4°C), and the cilia were pelleted with higher-speed centrifugation (17,000g, 30 min, 4°C). Cilia pellet was then resuspended in cilia final buffer (CFB) (50 mM HEPES pH 7.4, 3 mM MgSO4, 0.1 mM EGTA, 0.5% Trehalose, 1 mM DTT) supplemented with 1 mM PMSF. Sequential purification was performed to obtain doublets without protein structures bound outside. First, cilia membranes were removed from the axonemes by adding the final 1.5% Nonidet P-40 Alternative (Millipore, 492016) for 30 min. Then, the demembraned axoneme was spun down by centrifuge (7,800g, 10 min, 4°C). The pellet was resuspended with CFB supplemented with 1 mM PMSF and 0.4 mM ATP and incubated at room temperature for 5 min to release each doublet from the axoneme. After this, the final 0.6 M NaCl was added and incubated for 30 min on ice to deplete axonemal dyneins. The sample was centrifuged (16,000g, 10 min, 4°C) and resuspended in CFB containing 0.6 M NaCl to remove the remaining axonemal dyneins. Doublets were then dialyzed against low-salt buffer (5 mM HEPES pH 7.4, 1 mM DTT, 0.5 mM EDTA) at 4°C overnight to remove radial spokes. The clean doublets after dialysis were centrifuged again (16,000g, 10 min, 4°C), and the pellet was finally resuspended in B80-TK buffer (80 mM piperazine-N,N′-bis(2-ethanesulfonic acid) [PIPES] pH 6.9, 2 mM MgCl2, 1 mM EGTA, 1 mM DTT, 20 µM paclitaxel, 50 mM KCl).

### MT co-pelleting assay

Dynein-1 and dynein-2 MTBD constructs were ultracentrifuged (45,000 rpm for 30min at 4°C, himac S55A2 rotor) and the supernatant was used for the experiments. The concentrations of dynein-1 and dynein-2 MTBD constructs in the supernatant were determined based on the band intensities on SDS-PAGE gel using BSA as standards. 0.23 µM MTBD constructs were mixed with 120 µg/ml doublets in NAP5 buffer, incubated on ice for 45 min, and ultracentrifuged (45,000 rpm for 30min at 4°C, himac S55A2 rotor). The supernatant and pellet fractions were separated and analyzed by SDS-PAGE and the band intensities were analyzed.

### *In vitro* reconstitution

Purified doublets (final 500 μg/ml) were incubated with GST-Dyn2 (final 450 μg/ml) for 30 min at 25°C and then centrifuged (16,000g, 10 min, 4°C). The supernatant was removed, and the precipitation fraction was resuspended in B80-TK buffer and used for vitrification after adding BSA-treated gold nanoparticles (10-nm diameter) prepared as in (Iancu *et al*., 2006) at an optimal ratio.

### Cryo-ET analysis

3.5 μl sample was added to glow-discharged grids (Quantifoil R2/2) and vitrified using Vitrobot Mark IV (Thermo Fisher Scientific) with blot force 3 and blot time 3 sec. The grids were loaded into Titan Krios (Thermo Fisher Scientific), operating at 300 kV, equipped with Falcon II direct electron detector (Thermo Fisher Scientific). The tilt series were acquired from −40 to +60 degree tilts with 4-degree increments (26 images) with a total dose of 79.7 electrons. The nominal magnification was 29,000x, and the defocus was −0.5 or −1 μm. The phase plate’s activation time was 70 sec. The tomograms were reconstructed with Etomo from IMOD (Kremer *et al*., 1996) using the Simultaneous Iterative Reconstruction Technique (SIRT) method with binning of 3.

### Model building

Model building was performed as described in (Kubo & Bui, 2023). In brief, the tubulin lattice model for MD simulations was generated by MODELLER (Webb & Sali, 2016) using TUBA1B (UniProtKB: P68363) and TUBB (UniProtKB: P07437) sequences and cryo-EM structure of GDP MT (PDB ID: 6DPV) as the reference structure. For the detyrosinated tubulin lattice, the tyrosine residue at the C-terminus of the α-tubulin was removed. For the dynein-2 dimer model, the high-affinity model of the dynein-2 motor domain and the low-affinity model of the dynein-2 motor domain were generated as in (Kubo & Bui, 2023). The DYHC2 (UniProtKB: Q8NCM8) sequence was used for model building. For the low-affinity structure, dynein-2 motor domain structure in ADP·Vi condition (PDB ID: 4RH7) (Schmidt *et al*., 2015) was used as a reference. High-affinity dynein-2 model was generated by combining AAA+ and stalk region structure (PDB ID: 3VKH) (Kon *et al*., 2012), stalk and MTBD structure (PDB ID: 3J1T) (Redwine *et al*., 2012), and MTBD-tubulin interface (PDB ID: 6KIQ) (Nishida *et al*., 2020). First, 3VKH and 3J1T structures were combined by an MD simulations as in (Kubo *et al*., 2017). The model was combined with 6KIQ using COOT (Emsley & Cowtan, 2004). Finally, a homology model was generated using MODELLER (Webb & Sali, 2016).

### MD simulations

Four protofilaments each containing three tubulin dimers, and two dynein-2 motor domains corresponding to one dynein-2 dimer were used for the MD simulations. In each of the tyrosinated and detyrosinated tubulin lattice, the low-affinity dynein-2 head was placed at the center of the tubulin lattice ((1, 1) in Fig 4B), and the high-affinity dynein-2 head was placed in the PF on the far side of the paper (Fig 4A). The low-affinity dynein-2 structure was docked onto the tubulin lattice based on PDB ID: 3J1U (Redwine *et al*., 2012). High-affinity dynein-2 was placed in three positions relative to low-affinity dynein-2: backward, lateral, and forward (Fig 4A). The high-affinity dynein-2 head domain’s MTBD was set according to the MTBD position from PDB ID: 6KIQ (Nishida *et al*., 2020). 20 calculations were performed for each of the three initial positions as in (Kubo & Bui, 2023). MD simulations were performed by CafeMol version 2.1 (Kenzaki *et al*., 2011) for each simulation with 3 × 10^7^ MD steps. Underdamped Langevin dynamics at 323 K temperature were used for calculation. The friction coefficient was set to 2.0 (CafeMol unit), and default values were used for other parameters. For intra- and inter-molecules of the proteins, AICG2 + force field, electrostatic interaction, and excluded volume function were considered. Heat maps (Fig 4C and D) were generated based on the mass center-of-gravity coordinates of the MTBDs of the low-affinity dynein-2 head obtained from their trajectories.

### Data availability

The cryo-EM structures used for modeling are available from the Protein Data Bank under accession number 4RH7 for the low-affinity dynein-2; accession numbers 3VKH, 3J1T, 6KIQ for the high-affinity dynein-2; 3J1U for the initial position of low-affinity state dynein; 6DPV for tubulin lattice. All MD simulations in this paper were performed by CafeMol software (https://www.cafemol.org). The datasets analyzed in this study are available from the corresponding author upon reasonable request.

## Supporting information

Movie S1. Aligned-tilt series images of GST-Dyn2 decorated doublets collected with VPP.

Movie S2. A tomogram of GST-Dyn2 decorated doublets reconstructed from tilt series images in Movie S1.

## Acknowledgements

The authors appreciate Dr. Anthony J. Roberts from University of Oxford for generously providing GST-Dyn2 proteins and MTBD constructs. We thank Dr. Khanh-Huy Bui from McGill University for his insightful advice. Cryo-ET data with VPP was collected with help from the Facility for Electron Microscopy Research at McGill University, especially from Dr. Kaustuv Basu. This work was supported by Natural Science Foundation of Shanghai (24ZR1403800) and JST PRESTO (JPMJPR20E1) to MI, JSPS KAKENHI (22K15070) to SK, and the International Collaborative Research Program of Muroran Institute of Technology to AK. This preprint was typeset with the bioRxiv word template by @Chrelli: www.github.com/chrelli/bioRxiv-word-template

## Author contributions

MI conceived the project and designed the experiments. HKH and MI performed a culture of the cells and purification of the doublet with the help of AK. HKH and MI purified the MTBD constructs and performed biochemical analyses with the help of QHL. MI performed vitrification of the grids and collected data. MI and HKH performed the cryo-ET data analysis. SK performed the MD simulations. All authors were involved in the manuscript writing process.

---

**Figure S1.**
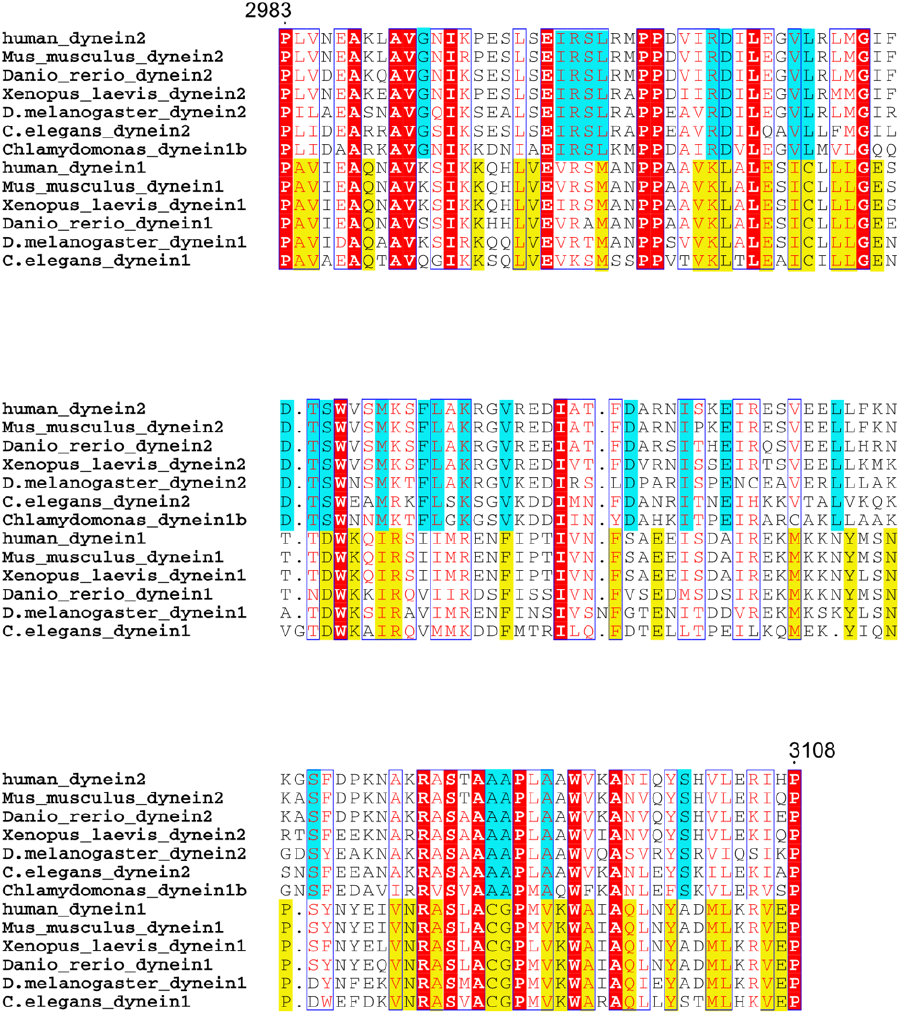
Sequence alignment of MTBD regions from dynein-2 and dynein-1 HCs. Amino acid sequence of the MTBD region of human dynein-2 HC (NCBI Reference Sequence: NP_001368.2), *Mus musculus* dynein-2 HC (NCBI Reference Sequence: NP_084127.2), *Danio rerio* dynein-2 HC (NCBI Reference Sequence: NP_001410228), *Xenopus laevis* dynein-2 HC (NCBI Reference Sequence: XP_041438615), *Drosophila melanogaster* dynein-2 HC (NCBI Reference Sequence: NP_001036369), *Caenorhabditis elegans* dynein-2 HC (NCBI Reference Sequence: NP_492221.2), *Chlamydomonas reinhardtii* dynein-1b HC (NCBI Reference Sequence: XP_001696428.1), human dynein-1 HC (NCBI Reference Sequence: NP_001367.2), *Mus musculus* dynein-1 HC (NCBI Reference Sequence: NP_084514.2), *Xenopus laevis* dynein-1 HC (NCBI Reference Sequence: XP_018086051.1), *Danio rerio* dynein-1 HC (NCBI Reference Sequence: NP_001036210.1), *Drosophila melanogaster* dynein-1 HC (NCBI Reference Sequence: NP_001261430.1), *Caenorhabditis elegans* dynein-1 HC (NCBI Reference Sequence: NP_491363.1) were aligned using Clustal W (Thompson, 1994) and figure was prepared using ESPript 3.0 (Robert & Gouet, 2014). Note that *Chlamydomonas reinhardtii* dynein-1b is IFT dynein equivalent to dynein-2. Cyan highlights show the amino acid residues conserved only in dynein-2 and yellow highlights indicate the amino acid residues conserved only in dynein-1.

**Figure S2.**
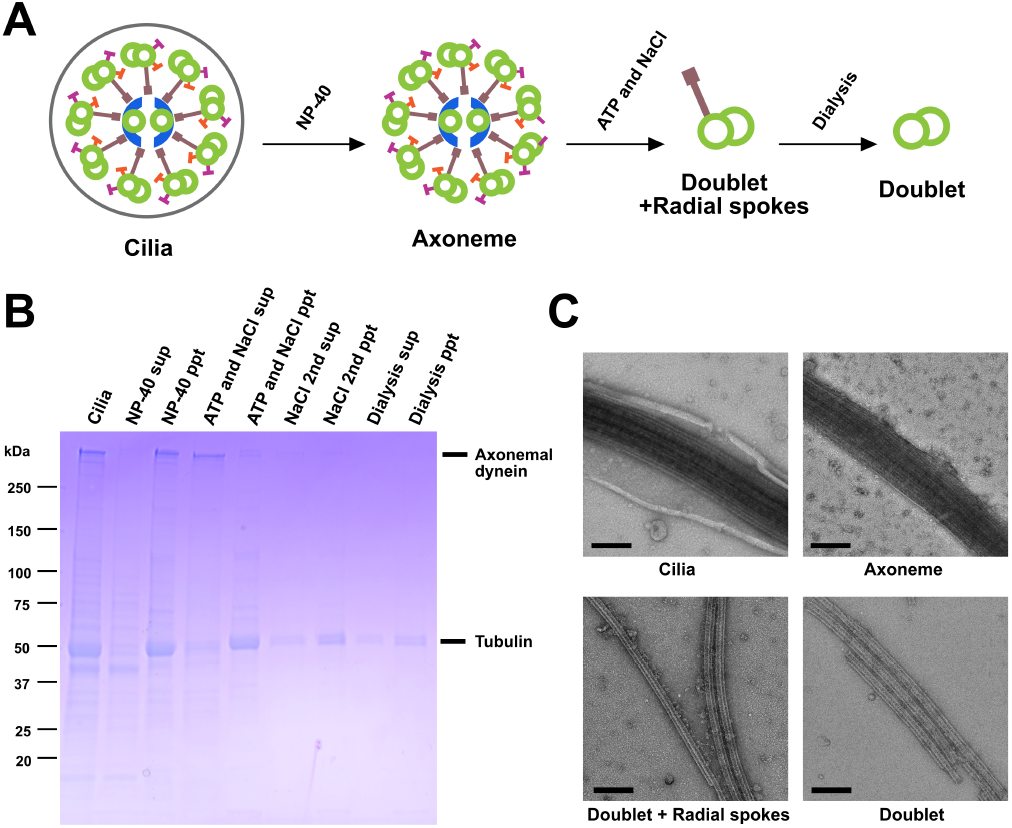
Preparation of doublets with clean outer surface. **(A)** The workflow of doublet purification with a clean outer surface. **(B)** SDS-PAGE gel of sequential purification of *Tetrahymena* doublet. Associated proteins, especially axonemal dyneins, are removed while the tubulin band is visible. **(C)** Negative staining EM images of sequentially purified doublet samples. The doublets with clean outer surfaces were obtained. Scale bars: 200 nm.

**Figure S3.**
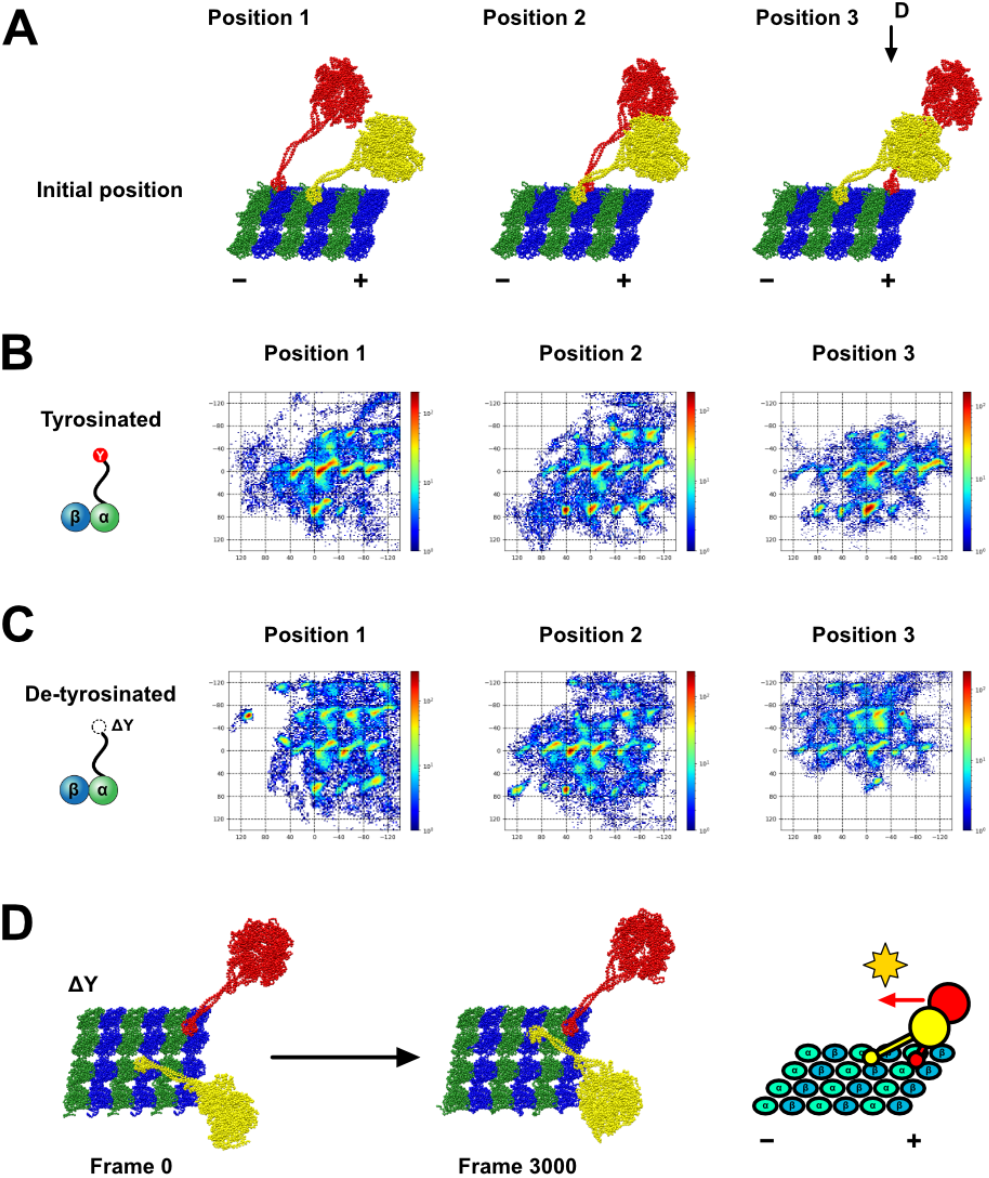
Data related to MD simulations of dynein2 dimer. **(A)** Initial structures of the MD simulations. Low-affinity dynein2 motor domain structure (yellow) was placed at the center of the tubulin lattice with high-affinity dynein2 (red) on the neighboring PF located at forward (position 1), adjacent (position 2), and backward (position 3). (−) and (+) indicate MT polarities. **(B-C)** Heat maps of the positions of the MTBD of low-affinity dynein-2 on tubulin lattice with tyrosinated tubulins (B) and detyrosinated tubulins (C). 20 trajectories are overlaid and colored depending on the frequencies. Blue color corresponds to low frequency and red color represents higher frequency. **(D)** Snapshots of MD simulation results showing that the low-affinity leading head is blocking the high-affinity trailing head of dynein-2 in the detyrosinated tubulin lattice. MD simulation was started from the initial state (frame 0, initial position 3 in (A)). In the simulated result (frame 3000), the low-affinity leading head moves in the direction of movement of the high-affinity trailing head. In the next cycle of the trailing head, the low-affinity leading head hinders the diffusional motion toward the minus end, and thereby, the dynein-2 dimer dissociates from the tubulin lattice (right panel). The view in (D) is indicated in (A).

**Movie S1. Aligned-tilt series images of GST-Dyn2 decorated doublets collected with VPP**.

**Movie S2. A tomogram of GST-Dyn2 decorated doublets reconstructed from tilt series images in Movie S1**.

